# EFFECT OF METHANOL STEM-BARK EXTRACT OF *Erythrina senegalensis* ON SERUM ALBUMIN LEVEL OF CARBON TETRACHLORIDE INDUCED LIVER DAMAGE IN WISTAR ALBINO RATS

**DOI:** 10.1101/2025.08.20.671200

**Authors:** Adetunji Abdulkabir, Raymond John, Turaki Aliyu, Tajudeen Yahaya, ThankGod Asekemhe, Azeem Kolawole

## Abstract

**Introduction:** Previous investigations already established the pharmacological importance of *Erythrina senegalensis*. The current study was conducted to determine the effect of methanol stem-bark extract of *Erythrina senegalensis* (MEES) on serum albumin level of carbon tetrachloride (CCl_4_) induced liver damage in wistar rats.

**Method:** Thirty wister rats of either sex weighing were randomly divided into six groups of five rats each. Group 1 served as the control received standard pelleted rat feed with normal distilled water. Group 2-6 were administered 30% CCl_4_ 1ml/kg body weight in liquid paraffin at 72hrs interval. Group 2 was untreated while group 3 was orally treated with standard drug salymarin for 14days. Group 4-6 were orally treated respectively with 100, 300 and 500mg/kg body weight of the MEES.

**Result:** The phytochemical analysis of the MEES indicated the presence of saponins, phenols, flavonoids, and tannins while the acute toxicity test showed no adverse reaction within the test population at a dose range of 200–2000mg/kg body weight. Unlike the control, inducement of CCL_4_ to Group2-6 indicated significant changes (p < 0.05) in the serum albumin level and hematological parameters. However, treatment with silymarin and increasing doses of the MEES restore the changes. The histopathological studies also reveal that increasing concentration of administered doses of the MEES significantly (p < 0.05) reduces cell necrosis around central vein.

**Conclusion:** The findings reveal that MEES at recommended doses can serve as alternative source for the development of safe, effective, and affordable liver-supplement drug.

## INTRODUCTION

Plant parts have been used to treat ailments since the dawn of time. There is historical evidence of the use of plants in the treatment of various illnesses throughout all civilizations (1). In the last decade, acute and chronic diseases have been treated with medicinal plants and over 80% of the world’s population depends solely on herbal plants as medicine (2). Plant material has remained useful sources of new drugs and becoming increasingly popular due to their perceived safety when compared to synthetic drugs. Plant remedies are also becoming more popular because they are less expensive, readily available, and simple to prepare and administer (1).

In Africa, the plant *Erythrinal senegalensis* is commonly employed by traditional healers for treatment of several ailments. In Northern Nigeria, the bark is used to treat jaundice and related diseases. In Ghana, the bark is recommended as emmenagogue and in French Guinea it is given to women after child birth (2). In Senegal the bark is considered as remedy for dysentery and colitis. In Mali, the flower, stem bark, leaves and roots are grinded separately and used for the treatment of amenorrhea, malaria, jaundice, diarrhea, and dysmenorrheal respectively (3). The pharmacological studies of this plant are demonstrated and include antibacterial, antiviral, antifungal, and antiplasmodial activities; it also induces analgesic and antipyretic action (2). As a result of common traditional usage and recorded pharmacological importance, the effect of the methanol stem-bark extract of *Erythrina senegalensis* (MEES) on serum albumin level of CCl_4_ induced liver damage in wistar rats was investigated.

## MATERIALS AND METHODS

### Experimental Animals

The approval of the Animal Ethics Committee of Federal University Birnin Kebbi, Nigeria, was sought before commencing this study. A total of forty two Wistar rats of either sex weighing between 89g and 179g were used for the study. They were purchased from the Animal House of Usman Danfodio University Sokoto, Sokoto state, Nigeria. The rats were acclimatized for one week prior to the commencement of the experiment, with 12-hours light/dark cycle maintained. The rats were fed with standard pelleted feed and potable drinking water, and were handled in accordance with the national and international ethical recommendations for care and use of laboratory animals.

### Sample Collection

*Erythrinal senegalensis* stem bark was collected on 4^th^ of August, 2021 from its natural habitat in Kango Wasagu, Danko Wasagu Local Government, Kebbi State, Nigeria. The collected stem bark of *E. senegalensis was* chopped into small sizes and air dried at room temperature sunlight to avoid denaturation of vital phytochemical constituents. The dry sample was crushed into small sizes using mortar and pestle and the obtained fine power form were stored in the freezer for further.

### Preparation of Methanol Extract

1.5kg of coarse powder of *E. senegalensis* in plastic rubber was macerated with 6L of methanol for 72hrs. The extract is decanted and allowed to stand, after which it is filtered through a Whatman No. 42 filter paper. The extract was concentrated using a rotary evaporator. The extract was transferred into a pre-weighed beaker and placed on a water bath at 600C until a plastic form of the extract is obtained. The crude extract was then weighed to determine the percentages yield of extraction. The percentage yield of extraction was calculated by using the following equation:

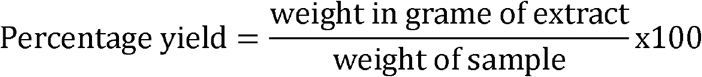

### Qualitative Phytochemical Screening

The screening for various phytochemicals present in the extract was carried out using standard methods as described below (2).

### Test for Saponins

0.2g of extract was shaken 5ml of distilled water in a test tube. Frothing which persists on warming indicates the presence of saponin.

### Test for Tannins

0.2g of the molten extract was mixed with few drops ferric chloride solution. A blue-black, green or blue-green precipitate indicates the presence of tannin.

### Test for Alkaloid

0.2g of the molten extract was mixed with little amount of HCl and then wagners reagent. Formation of a white precipitate indicates the presence of alkaloid.

### Test for Phenols

0.2g of molten extract was mixed with ferric chloride solution. A green or dirty green precipitate indicates the presence of phenol.

### Test for Terpenoid

About 0.2g extract was mixed with 2ml Chloroform and 3ml of concentrated sulphuric acid was added carefully to form a layer. A reddish brown coloration of the interface formed indicates the presence of terpenoids

### Test for Flavonoid

0.2g of the molten extract was mixed with 1ml of 2% ammonium chloride and the exposed to light. Yellow precipitate indicates the presence of Flavonoid.

### Quantitative Phytochemicals Dertermination

The quantitative determination of detected phytochemicals in the extract was carried out using standard methods as described below (4).

### Determination of Total Tannins

5g of the extract was weighed into a 100-ml plastic bottle containing 50 ml of distilled water and shaken for 1hr in a mechanical shaker. The filtrate obtained was put in a 50-ml volumetric flask and made up to the mark. Then, 5 ml of the filtrate was pipette into a tube and mixed with 3 ml of 0.1M FeCl3 in 0.1 N HCl and 0.008M potassium ferrocyanide. The absorbance was measured in a spectrophotometer at 120 nm wavelength within 10 min. A blank sample was prepared, and the color also developed and read at the same wavelength. A standard was prepared using tannic acid to get 100 ppm and measured. The weight was thereafter expressed as a percentage of the raw extract.

### Determination of Saponin

20g of the extract was transferred into 200 ml of 20% ethanol. The suspension produced was heated over a hot water bath for 4hr with continuous stirring at about 55 °C. The mixture was filtered, and the residue re-extracted with another 200 ml of 20% ethanol. The combined extracts were reduced to 40 ml over a water bath at about 90 °C. The concentrate was transferred into a 250-ml separator funnel, and 20 ml of diethyl ether was added and shaken vigorously. The aqueous layer was recovered while the ether layer was discarded. The purification process was repeated, and 60 ml of n-butanol was added. The combined n-butanol extracts were washed twice with 10 ml of 5% aqueous sodium chloride. The remaining solution was heated in a water bath. After evaporation, the samples were dried in the oven to a constant weight and estimated as a percentage of the raw extract.

### Determination of Phenolic content

The assay for determination of total phenolic contents was carried out using Folin - Ciocalteau method. 0.5g of gallic acid was dissolved in 50ml of ethanol. Then, 6.5mL of distilled water was added to 0.5 mL of the extract in test-tube. 0.5 mL of a tenfold diluted (2 mL, 1:10 diluted with distilled water) Folin Ciocalteau reagent was added and 7.5% sodium carbonate was then added. The mixture was allowed to stand for 90 min at room temperature; absorbance of the solutions was measured at 729nm using spectrophotometer. All determinations were performed in triplicates with Gallic acid utilized as the positive control. The total phenolic contents were expressed as mg of Gallic acid equivalents (GAE) per g of dried extracts.

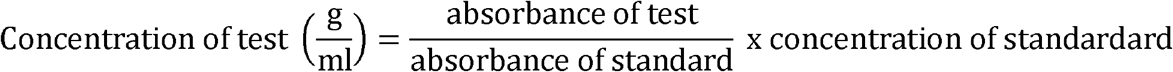

### Determination of Flavonoids

A sample solution was prepared by dissolving 0.05g of the sample extract in a conical flask and was filled up to 50mls. 2ml of 2% Alcl_3_ was dispensed into the three sets of test tube containing 2ml of extract and also into three empty test-tubes. 2mls of the standard (Quercetin) was measured into test tubes containing only AlCl_3_ and shaken, and cover and was also properly. The solutions were incubated for one hour (60 minutes) at room temperature and absorbance was taken at 420nm.

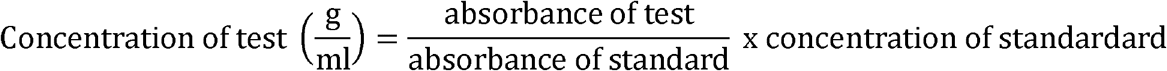

### Determination of 2,2-Diphenyl-1-Picrylhydrazyl (DPPH) Radical Scavenging Assay

A solution of 0.1 mM DPPH in methanol was prepared, and 0.5ml ≈500µL of this solution was mixed with 2mL of extract in methanol at different concentrations (25, 50, 75 and 150 µg/ml). The experiment was repeated three times at each concentration. The reaction mixture was vortexed thoroughly and left in the dark for 30 min. The absorbance of the mixture was measured spectrophotometrically at 517 nm (5).

Percentage DPPH radical scavenging activity was calculated by the following equation:

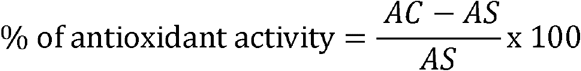

Where: AC is the absorbance of the control and

AS is the absorbance of the extracts (samples).

### Acute toxicity (LD50) Study

The acute toxicity study of the stem bark methanol extract of *E. senegalensis* was determined using modified OECD method (6).

A total of 12 wistar rats, divided into 4 groups. The first group received the extract at a dose of 2000 mg/kg body weight; second group received the extract at a dose of 1000 mg/kg body weight, third group served as the control while the last group received the extract at the dose of 200 mg/kg body weight. Animals are observed individually after dosing at least once during the first 30 minutes, periodically during the first 24 hours, with special attention given during the first 4 hours and daily thereafter. The animals were observed for mortality, body weight, clinical signs and gross pathological changes through day 14 and the weight of the rat is weighed weekly.

### Experimental Design of Hepato-Protective Studies

A total of 30 wistar rats were used. The rats were grouped according to similar body weights, into 6 groups of five rats each and treated orally as follows for 14 days, 72hrs after the induction of liver damage:

**Table 1.**
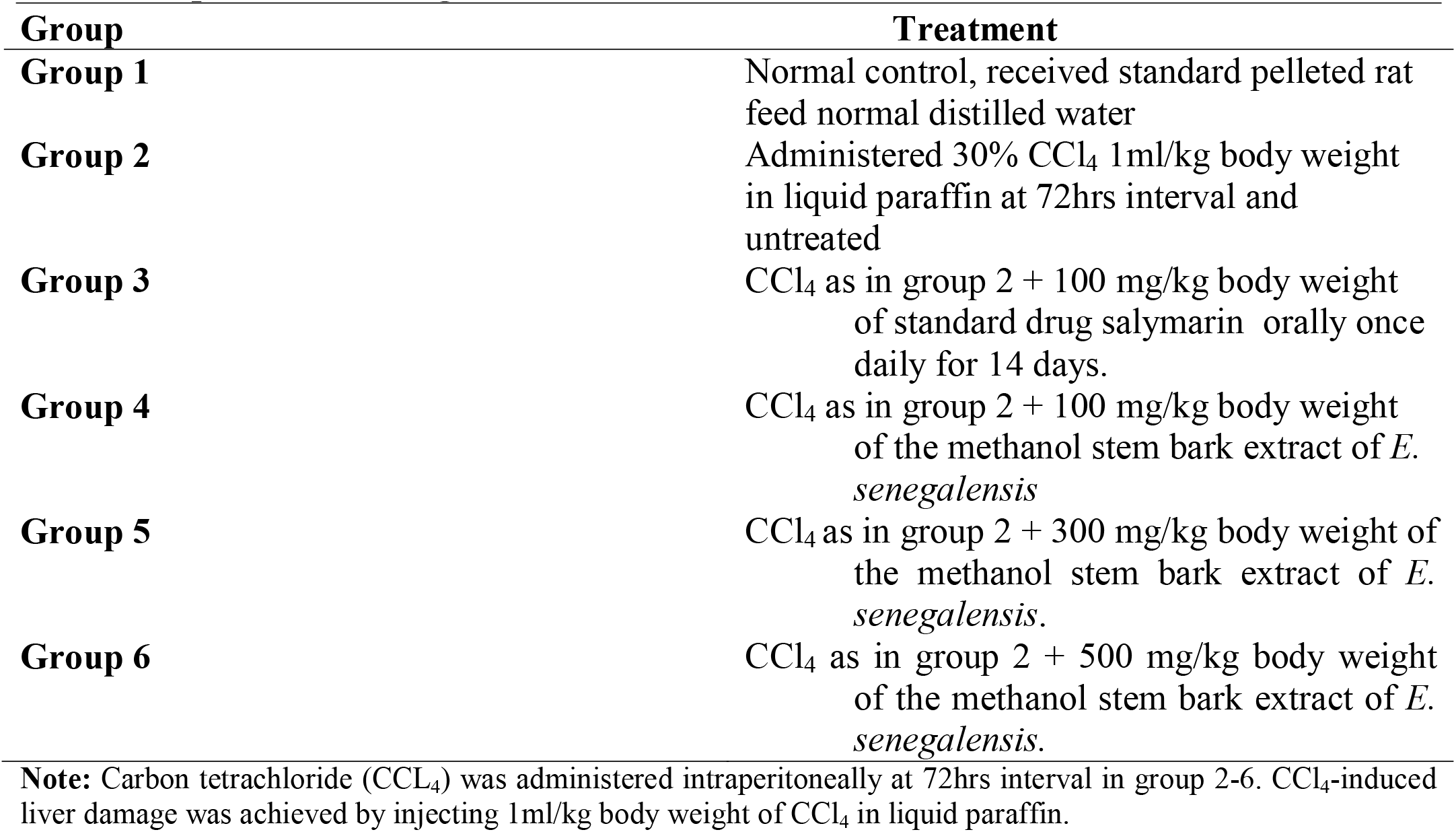
Experimental Design.

### Blood and Tissue Collection

The procedures of Joshua et al. (7) were followed in blood collection and preparation prior to analysis. At the end of the 14 days, the overnight fasted rats were sacrificed under chloroform anesthesia. Blood samples were collected by cardiac puncture into Ethylene Diamine Tetra-acetic Acid (EDTA) and sterile plain bottles for haematological and serum biochemical studies, respectively. The abdominal cavities of the rats were opened to excise the liver for histological examinations.

- **Hematological Studies** The EDTA blood samples were used for hematological investigations using an automated hematological analyzer (Sysmex KX-21, Japan) for the measurement of hemoglobin (Hb) levels, platelet count (PLC), corpuscular hemoglobin (MCH),total count of red blood cells (RBC), and white blood cells (WBC) (7).
- **Estimation of Serum Albumin** The blood samples in the sterile plain bottles were allowed to clot and serum was prepared by centrifugation at 3000 rpm for 15 minutes. The albumin level is measured using a dye-binding method, where the addition of bromcresol green dye to the serum forms a stable complex, allowing for accurate quantification. The absorbance of the samples and of the standard was measured against reagent blank at 546 nm, and temperature of 37°C. These tubes and their contents were mixed and incubated for 90 minutes at 37°C. Estimation of albumin level (g/dl) was obtained using a spectrophotometer (7).
- **Histological Examination** Harvested organs were trimmed of adherent tissues and cleaned using physiological saline solution. The organs were weighed and fixated in 10% neutral buffered formalin. The tissues were embedded in paraffin wax, sectioned to a thickness of 5 µm and stained using Hematoxylin and eosin dyes. Tissues were mounted on glass slides and examined under a light microscope for inflammations, scarring, and fibrosis (7).

### Statistical Analysis

The data obtained were analyzed using excel software. Tests of statistical significance were carried out using the one-way analysis of variance (ANOVA). The results were expressed as mean ± standard deviation. p values < 0.05 were considered statistically significant.

## RESULTS

### Qualitative screening of the extract

Table 2 shows the results of qualitative analysis of the MEES. The phytochemicals detected are tannins, phenols, saponins and flavonoids.

**Table 2.**
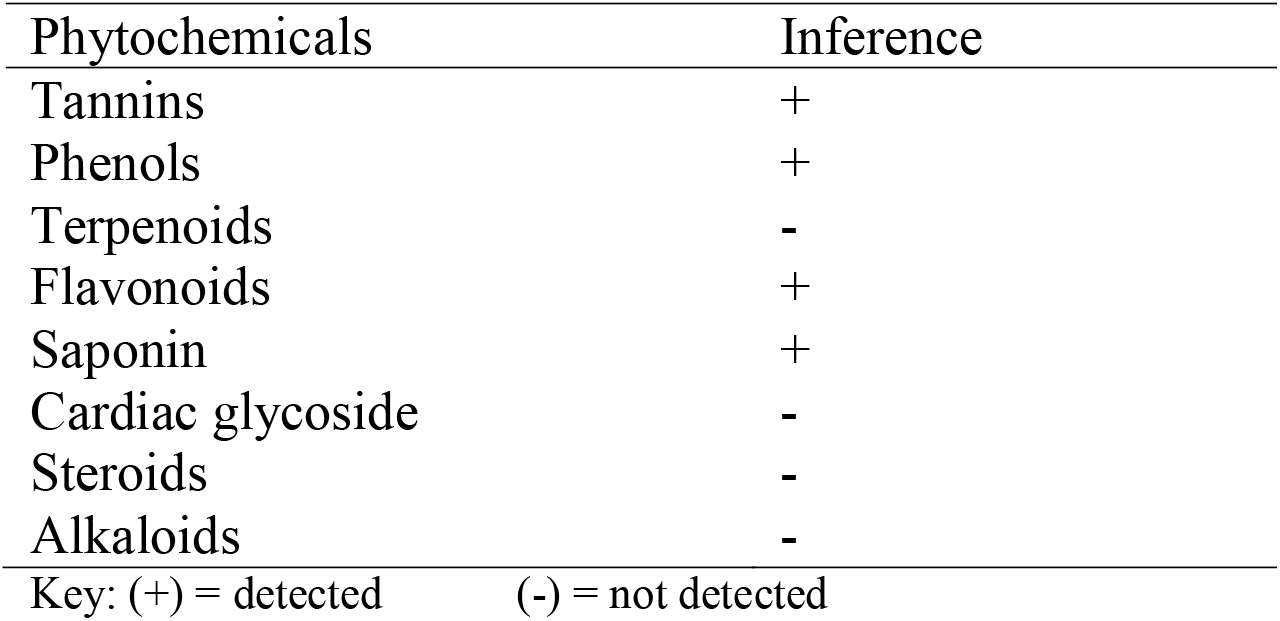
Phytochemical Constituents of Methanol Stem-bark Extract of *Erythrina senegalensis*.

| Phytochemicals | Inference |
| --- | --- |
| Tannins | + |
| Phenols | + |
| Terpenoids | - |
| Flavonoids | + |
| Saponin | + |
| Cardiac glycoside | - |
| Steroids | - |
| Alkaloids | - |

Key: (+) = detected (-) = not detected

### Qualitative value of the detected phytochemical consitutents

The values of the detected phytoconstituents are presented in Table 3. Flavonoids are the most abundance followed by phenols, tannins and saponins.

**Table 3.**
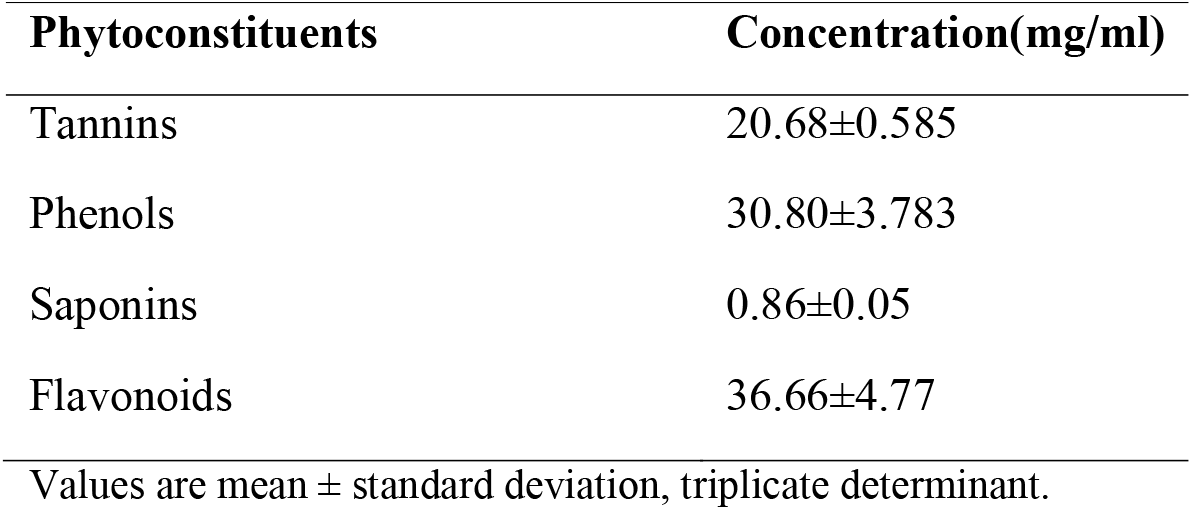
Quantitative value of detected phytoconstituents.

| Phytoconstituents | Concentration(mg/ml) |
| --- | --- |
| Tannins | 20.68±0.585 |
| Phenols | 30.80±3.783 |
| Saponins | 0.86±0.05 |
| Flavonoids | 36.66±4.77 |

Values are mean ± standard deviation, triplicate determinant.

### In-Vitro Antioxidant Activity Using DPPH Radical Scavenging Assay

Table 4 shows the result of the antioxidant activities of the MESS using DPPH radical scavenging assay. The antioxidant activity of the MEES increase sequentially with increase in the MEES concentration

**Table 4.** Percentage Inhibition of DPPH Radical by Methanol Stem Bark Extract.

| Conc. (µg/ml) | Antioxidant activity (%) |
| --- | --- |
| 25 | 58.40 |
| 50 | 62.45 |
| 75 | 66.20 |
| 150 | 72.40 |

### Acute toxicity (LD50) of the MEES

The acute toxicity test of the MEES showed no mortality or adverse reaction within the test population at a dose range of 100–2000mg/kg body weight. This showed the relative safety of the MEES. Table 5 summarizes the acute toxicity of the MEES.

**Table 5.** Lethal Dose (LD50) of the Methanol Stem-Bark Extract of *E. senegalensis*.

| Group | Dosage(mg/kg body weight) | Mortality(D/T) | Symptoms | Difference in mean weight (g) (final – initial) |
| --- | --- | --- | --- | --- |
| Group 1 | 2000 | 0/3 | None | 217 – 192 = 25 ↑ |
| Group 2 | 1000 | 0/3 | None | 129 – 100 = 29 ↑ |
| Group 3 | 0 | 0/3 | None | 180 – 151 = 29 ↑ |
| Group 4 | 200 | 0/3 | None | 198 – 174 = 24 ↑ |

### Hepatoprotective activity of the MEES on serum albumin level of CCl_4_ induced liver damage of wistar rats

Table 6 shows the hepatoprotective activity of the MEES on the CCL_4_ induced liver damage of the wister rats. Increase in concentration of the administered MEES leads to sequential increase in the value of the serum albumin, thereby bringing the value closer to the serum albumin level of the salymarin drug administered group and the control group. Figure 1 summarizes the value of the serum albumin level of each group.

**Table 6.**
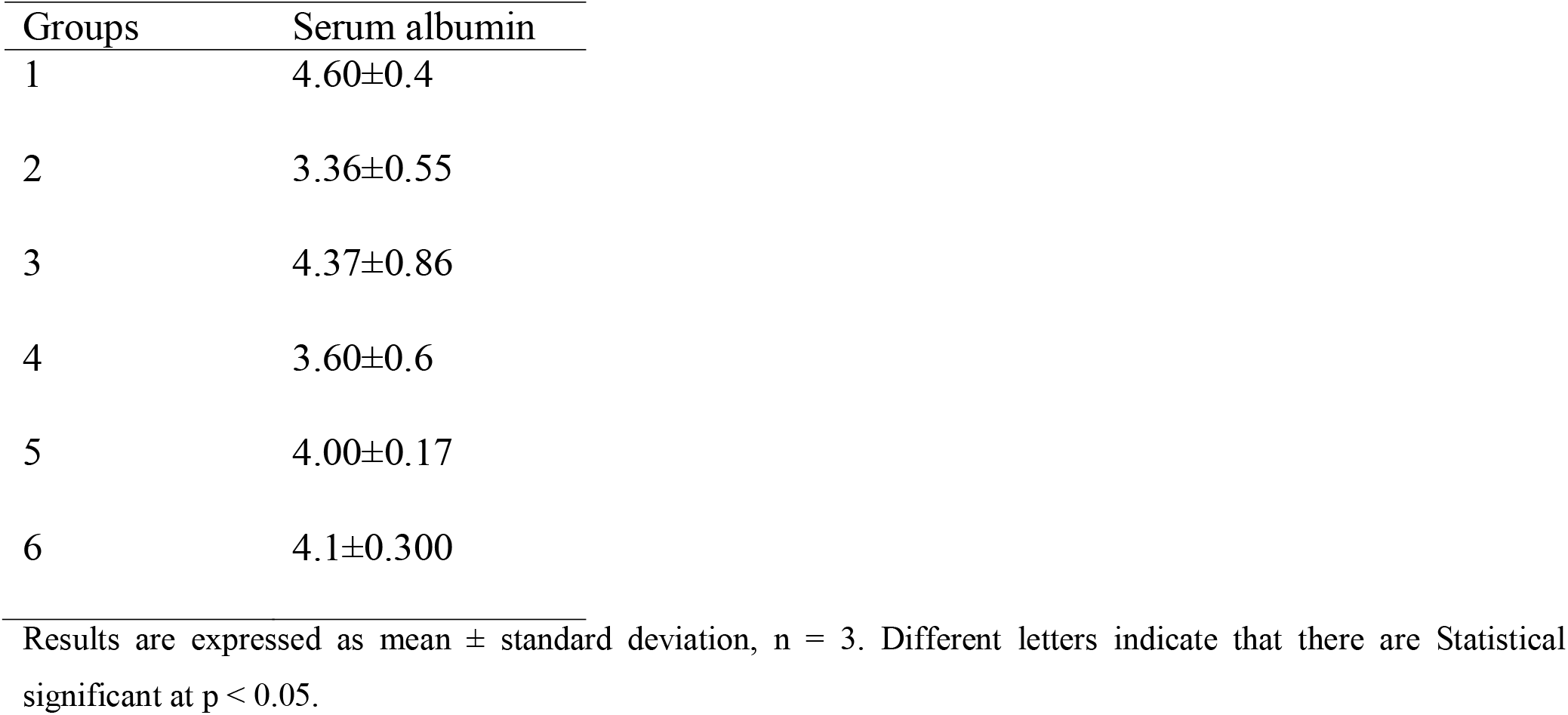
Serum Albumin Level of CCl_4_ Induced Liver Damage of Wistar Rats Treated with MEES and Salymarin.

**Figure 1.**
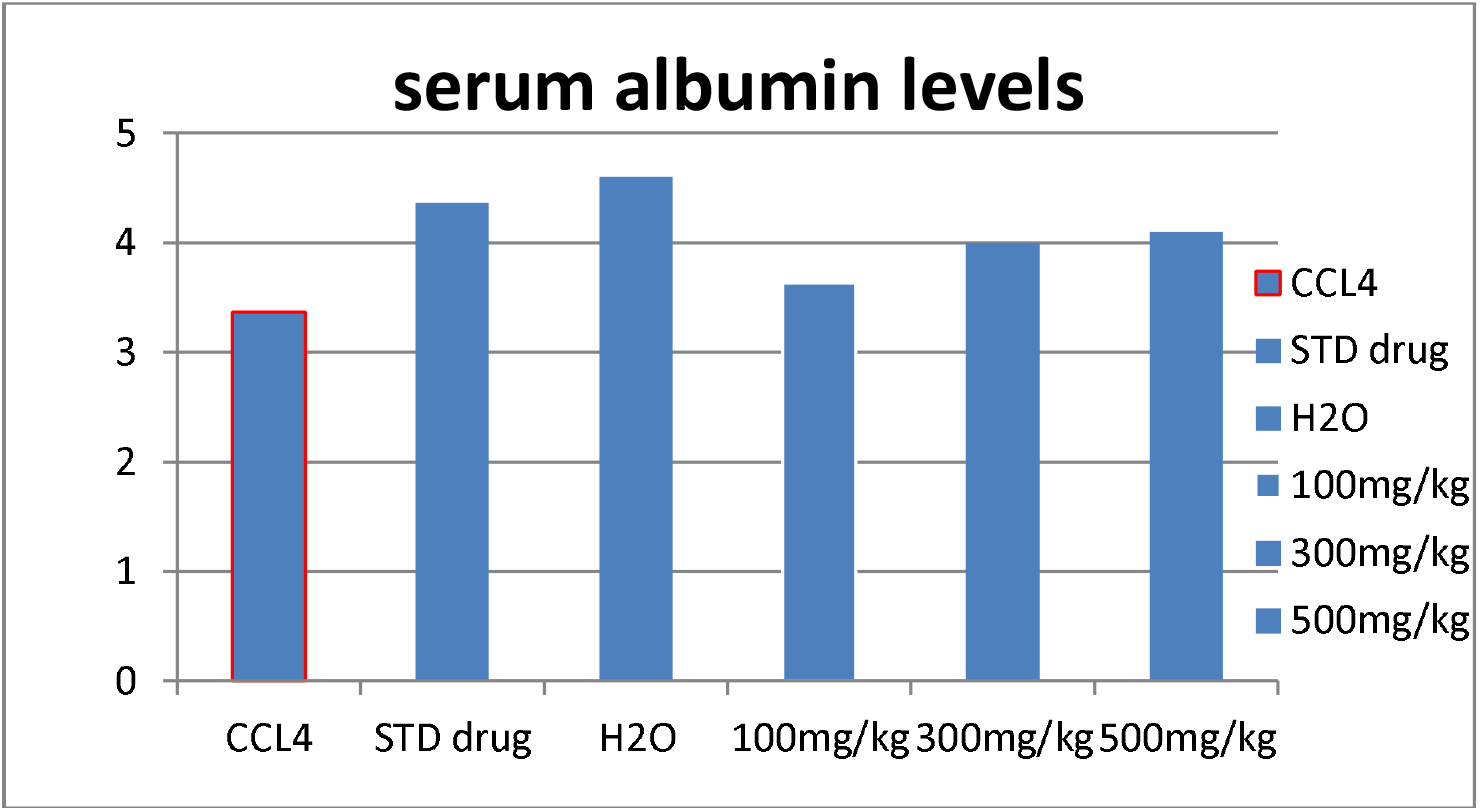
Mean Distribution of serum albumin (g/dl) within Experimental Animal Group

### Hematological Profile of MEES treated CCl_4_ Induced Liver Damage of Wistar Rats

Table 7 shows the values of the hematological investigation carried out on the CCl_4_ induced liver damage of the wisters treated with the MEES. The RBC count concentrations of the negative control induced-CCl_4_ untreated group were observed to be lower when compared to the normal control. Treatment with silymarin and increasing doses of the MEES indicated increase in the RBC count which is comparable to that of the normal control. The WBC, Hb, MCV and PLT count of the induced-CCl_4_untreated group was observed to be significantly higher when compared to the normal control. However, Treatment with silymarin and increasing doses of the methanol stem bark of *E. senegalensis* recorded a significantly lower of these parameters when compared to the untreated group.

**Table 7.** Effect of the Methanol Stem Bark of *Erythrina senegalensis* on Haematological Profile of CCl_4_ Induced Liver Damage in Wistar Rats.

| Groups | WBC count (x 10<sup>9</sup>cells/L) | RBC count (×10<sup>12</sup> cells/L) | HB conc. (g/dl) | MCH (pg) | PLT count (x10<sup>9</sup>/L) |
| --- | --- | --- | --- | --- | --- |
| 1 | 0.86 ±0.38<sup>a</sup> | 5.99±1.61<sup>a</sup> | 8.75±2.05<sup>a</sup> | 14.60±5.8a | 244 ± 2.83<sup>b</sup> |
| 2 | 2.2±0.28<sup>b</sup> | 4.46±0.59<sup>c</sup> | 12.95±0.49<sup>ac</sup> | 29.03±6.7 | 368 ± 2.83<sup>ac</sup> |
| 3 | 1.03±0.29<sup>c</sup> | 5.55±1.68<sup>a</sup> | 10.95±3.74<sup>c</sup> | 19.72±4.86c | 297±1.41<sup>a</sup> |
| 4 | 1.05±0.07<sup>c</sup> | 4.92±0.42b | 11.43±0.44<sup>ac</sup> | 23.23±0.28 | 265±1.41 |
| 5 | 0.71±0. 55<sup>ac</sup> | 4.94±1.33<sup>b</sup> | 9.1±0.14<sup>b</sup> | 18.42±5.3b | 205±2.83 |
| 6 | 0.65±0. 64<sup>ac</sup> | 5.47±0.66<sup>a</sup> | 8.7±2.97<sup>a</sup> | 15.90±6.15 | 292±5.66<sup>a</sup> |

Results are expressed as mean ± standard deviation, n = 3. Different letters indicate that there are Statistical significant at p < 0.05. WBC = white blood cell; RBC = red blood cell; HB = hemoglobin; MCH = mean corpuscular hemoglobin; PLT = plate

### Histopathological studies of the MEES on CCl_4_ Induced Liver Damage in Wistar Rats

Figure 2 describe the effect of the MEES on the liver tissue of each group. **G1** is the liver section of normal control rats, showing a normal hepatic architecture represented by hepatic lobule with a thin walled central vein. **G2** is the liver section of untreated induced CCL_4_ rats, showing cell necrosis around Central vein (CV) with inflammatory cells and fibroblasts, and degenerating hepatocytes in the portal area. **G3** is the liver section of induced CCl_4_ rats treated with silymarin, showing disappearance of fat droplets from hepatocyte cytoplasm, regeneration of the central vein, hydropic degeneration in the hepatocytes, and regeneration of hepatic parenchyma. **G4** is the liver section of induced CCl_4_ rats treated with 100mg/kg, showing degenerating hepatocytes in the portal area, collapsed of CV and inflammatory cells. **G5** is the liver section of induced CCl_4_ rats treated with 300mg/kg show restoration of CV and regeneration of cells. G6 is the liver section of induced CCl_4_ rats treated with 500mg/kg showing repair of the liver such as restoration of CV and regeneration of hepatocytes.

**Figure 2.**
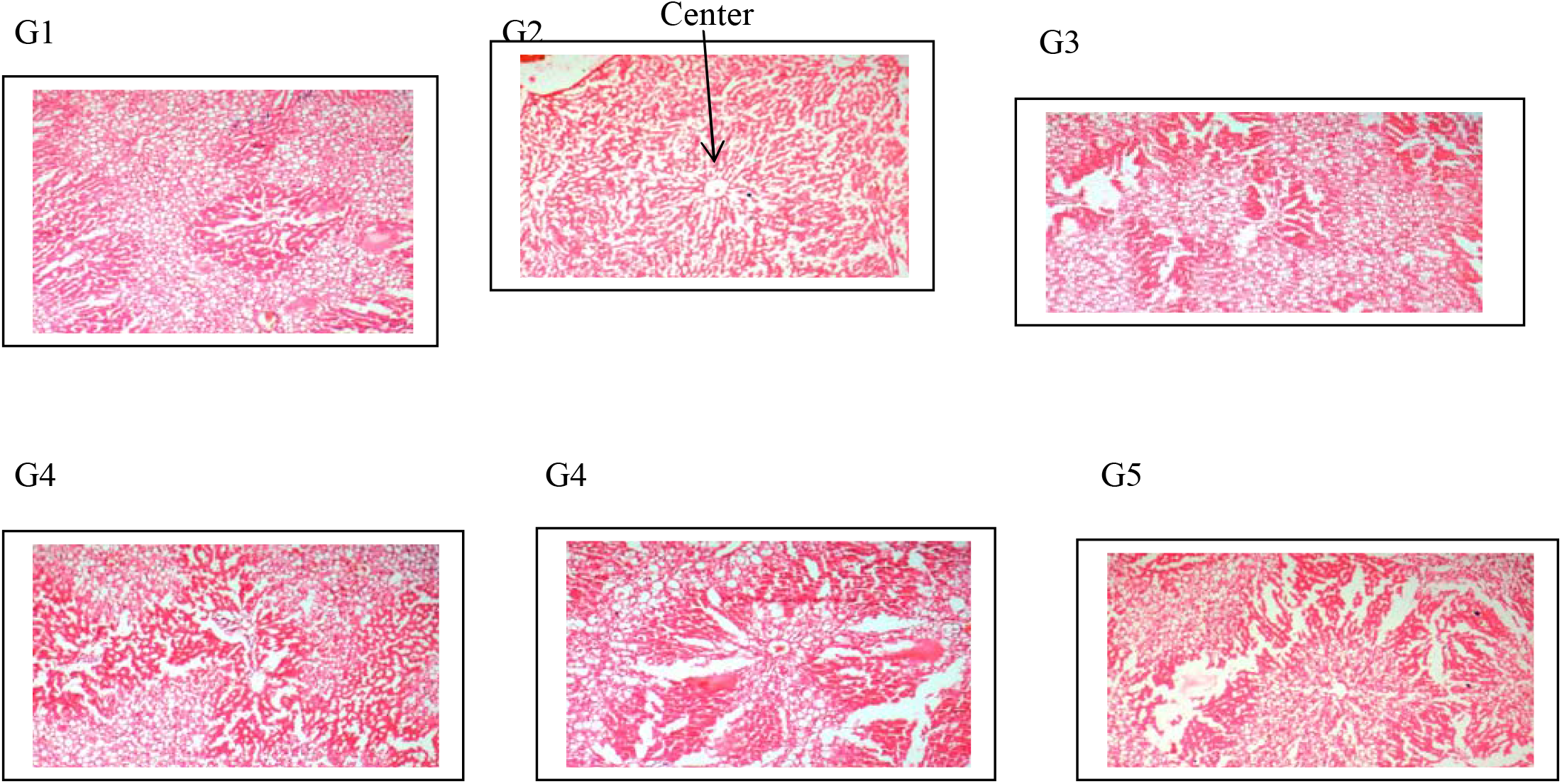
**G1**: Photomicrograph of the liver of normal control rats group. G2: Photomicrograph of the liver induced-CCl_4_ untreated rats group. G3: Photomicrograph of the liver of the negative control rats group. G4: Photomicrograph of the liver of the rat group treated with 100mg/kg of the MEES. G5: Photomicrograph of the liver of the rat group treated with 300mg/kg of the MEES. G6: Photomicrograph of the liver of the rat group treated with 500mg/kg of the MEES.

## DISCUSSION

The study was carried out to determine the effect of methanol stem-bark extract of *Erythrina senegalensis* (MEES) on serum albumin level of CCl_4_ induced liver damage in wistar rats. The MEES was discovered to contain phytochemicals which include saponin, phenols, flavonoids, and tannins. These phtochemicals have been reported to have beneficial effects to human health (8). The quantitative determination of the phytoconstituents is similar to the report of Parker et al. (9). The result obtained could explain why stem bark of *Erythrina senegalensis* is widely used in Africa to treat GIT related diseases (3).

Acute toxicity (LD50) of the methanol stem-bark extracts of *Erythrina senegalensis* (MEES) showed that the MEES did not cause any physical sign of toxicity, behavioral changes or mortality. This indicates that the plant is safe for consumption and for medicinal purposes. Following administration of CCL_4_, changes such as weight loss, dull fur, yellowing of the eyes and reduction in activities was observed among the rats in the test groups. Treatment with increasing dosage of the MEES led to reduction in the highlighted physical changes among the test groups. The hematological studies show that increase in concentration of administered MEES significantly (p<0.05) increase the RBC levels of the treated test groups when compared with the untreated test group. This result is similar to investigation carried out by Parker et al. (9), whose discovery linked low RBC level to abnormal hemolysis and liver cirrhosis, and increase in RBC level of the treated group to liver restorative effect of extract of *Erythrina senegalensis* (9). The WBC, Hb, MCH and PLT count of the untreated test group was observed to be significantly higher (p<0.05) when compared to the treated test groups and control group. Elevated WBC, Hb, MCH and PLT count indicate inflammatory response to CCL4-induced liver damage (10).

Liver proteins and enzymes are used to evaluate liver function (9). The level of the protein-serum albumin is commonly used as marker of liver function. In this study, the serum albumin level of the untreated test group is extremely low compared to the control group and treated groups, indicating hepatocellular damage. Increasing the concentration of the MEES significantly (p<0.05) increase the serum albumin level, signifying hepatoprotective effect of the MEES. The result agreed with the report of Arunsi et al. (11), who reported that plant extracts rich in flavonoids, tannins and phenols, do exhibit hepatoprotective properties (11). Phytochemically, the methanol stem-bark extract of *Erythrina senegalensis* contains required health percentage of these phtoconstituents indicating its hepatoprotective roles in CCl_4_ induced liver damage.

## CONCLUSION

The study has shown that the methanol stem-bark extract of *Erythrina senegalensis* contains beneficial phytocontituents like tannins, phenols, saponins and flavonoids, which exhibit hepatoprotective properties. In the light of our finding, the MEES exhibited these hepatoprotective effects by improving the serum albumin level and normalizing the hematological profile of CCl_4_-induced liver damage of wister rats. This show that methanol stem-bark extract of *Erythrina senegalensis* can serve as alternative source for the development of safe, effective, and affordable liver supplement.

## RECCOMENDATION

More studies should be carried out to verify these claims with other parts of the plant, and the level of the plant toxicity should be further determined with higher concentration of extract.

## CONFLICT OF INTERES

The authors declare no conflict of interest.

